# CALCIUM-DEPENDENT PROTEIN KINASE32 regulates cellulose biosynthesis through post-translational modification of cellulose synthase

**DOI:** 10.1101/2023.02.01.526621

**Authors:** Xiaoran Xin, Donghui Wei, Lei Lei, Haiyan Zheng, Ian S. Wallace, Shundai Li, Ying Gu

**Author notes:** Present address: Nature Plants, One New York Plaza, Suite 4500, New York, NY 10004 USA. These authors contributed equally to this work (Xiaoran Xin, Donghui Wei).

## Abstract

Cellulose is an economically important source of food, paper, textiles, and biofuel. As an essential component of plant cell walls, cellulose is critical for plant cell growth. Despite its economic and biological significance, the regulation of cellulose biosynthesis is poorly understood. Phosphorylation and dephosphorylation of cellulose synthases (CESAs) were shown to impact the direction and velocity of cellulose synthase complexes (CSCs). Despite a high prevalence of phosphorylation sites in CESAs, the protein kinases that phosphorylate CESAs are largely unknown. Here, we demonstrate that CALCIUM-DEPENDENT PROTEIN KINASE32 (CPK32) regulates cellulose biosynthesis *via* phosphorylation of CESA3. Phosphorylation of CESA3 is important for the motility and stability of CSCs. Hence, we uncovered a new function of CPKs that regulates cellulose biosynthesis and a novel mechanism by which phosphorylation regulates the stability of CSCs.

## Introduction

Reversible protein phosphorylation plays a critical role in diverse biological processes in eukaryotes. Most common phosphorylation or dephosphorylation events occur on serine, threonine, and tyrosine residues, while less stable phosphorylation modifications also occur on other residues (1). Phosphoproteomic studies revealed that more than 67% of human proteome and 47% of *Arabidopsis* proteome were phosphorylated, contributing to over 200,000 phosphorylation sites in human and 43,000 phosphorylation sites in *Arabidopsis*. These phosphorylation sites are modified by 518 kinases / 226 phosphatases in human and over 1000 kinases / 112 phosphatases in *Arabidopsis* (2–4). The regulation of proteins via phosphorylation and dephosphorylation can be extremely complex. For example, the regulation of mitophagy is mediated by different phosphoforms of Dynamin-related Protein 1 (DRP1) in mammalian cells. The phosphorylation of DRP1 on Ser^40^, Ser^44^, Ser^585^, and Ser^616^ promote mitochondrial fragmentation and subsequent mitophagy (5–7). Mitochondrial fusion, on the other hand, is promoted by DRP1 phosphorylation of Ser^637^ and dephosphorylation on Ser^616^ (8, 9). These modifications are mediated by different kinases and phosphatases under various cellular conditions (10). Another elegant example is the regulation of immune response in *Arabidopsis* through the degradation of BOTRYTIS-INDUCED KINASE1 (BIK1), which serves as the signaling hub linking Pattern Recognition Receptors (PRRs) complex and downstream effectors (11–13). Phosphorylation of CALCIUM-DEPENDENT PROTEIN KINASE 28 (CPK28) on Thr^76^ and Ser^318^ mediated by itself and BIK1 activates its phosphorylation of two E3 ligases, which induce BIK1 polyubiquitination and immune response. Phosphorylation of these sites is also required for CPK28 ubiquitination and degradation mediated by ARABIDOPSIS TOXICOS EN LEVADURA 31/6, which resists overactivation of immune signaling (14–16). Thus, it indicates a fine-tune immune signaling regulation via the phosphorylation statues of CPK28.

Cellulose, as the major load-bearing structure in plant cell walls, is the most abundant feedstock for sustainable products on earth. Despite its importance, there are limited studies of the regulation of cellulose synthesis. In higher plants, cellulose is synthesized by plasma membrane-localized cellulose synthases (CESAs) (17). Genetic studies have revealed that different CESA isoforms are responsible for cellulose synthesis in different types of cell walls. CESA1, −3, −6, and −6-like (CESA2, −5, and −9) are involved in primary cell wall synthesis, whereas CESA4, −7, and −8 are required for secondary cell wall formation (18–20). So far, two types of post-translational modification of CESAs have been empirically demonstrated: S-acylation and phosphorylation. S-acylation is important for CESA trafficking from Golgi to the plasma membrane as mutation of cysteine residues in CESA7 failed to localize to the plasma membrane (21, 22). Phosphoproteomic analyses revealed that most of the phosphorylation events occur at the N-terminal hypervariable regions of CESAs and a few phosphorylation sites reside in the catalytic domains of CESAs (19, 23–25). The physiological impact of these phosphorylation sites were examined by expressing the mutated version of CESA, e.g. serine (S) and threonine (T) to Alaine (A) that eliminates phosphorylation or glutamine (E) that mimic the phosphorylation, in the corresponding CESA knock-out or knock-down background (26–28). For instance, phospho-dead and phosphomimic mutations of CESA1 differently affected *Arabidopsis* hypocotyl elongation, root length, and CSC moving directionality (27). A similar study was carried out to assess the impact of CESA3 phosphorylation, where the phosphorylation and dephosphorylation of Ser^211^ and Thr^212^ at the N-terminal region of CESA3 differently impacted anisotropic cell expansion, cell wall composition, and CSC bidirectional motility (26). Furthermore, a red/far-red light photoreceptor PHYTOCHROME B (PHYB) has been demonstrated to regulate cellulose synthesis via impacting the phosphorylation status of CESA5 (28). Activation of PHYB is sufficient to restore CSC motility in *cesa6* mutant, which is phenocopied by introducing the phosphomimic CESA5 in *cesa6* mutant (28). Evidence from these studies suggests that the phosphorylation plays a critical role in the regulation of cellulose synthesis.

Despite a high prevalence of phosphorylation sites in CESAs, there is limited characterization of kinases that mediate those phosphorylation events. BRASSINOSTEROID INSENSITIVE2 (BIN2) was shown to phosphorylate a peptide derived from *Arabidopsis* CESA1 containing Thr^157^ (29). BIN2 is a glycogen synthase kinase 3-like kinase, which acts as a negative regulator in brassinosteroid signaling (30, 31). The phosphorylation of CESA1 by BIN2 negatively impacts CSC motility (29). A *BIN2* gain-of function mutant, *bin2-1*, exhibits reduced cellulose content and CSC motility, both of which are restored by the application of a BIN2 inhibitor, bikinin (29). Considering the numerous pairwise combinations of phosphorylation sites and kinases *in vivo*, the kinases that catalyze the phosphorylation of CESAs are most likely not limited to BIN2.

To identify additional kinases that phosphorylate CESA, a yeast two-hybrid screening was performed using the catalytic domain of CESA3 (CESA3CD) as the bait to search for potential CESA3-interactive partners. CALCIUM-DEPENDENT PROTEIN KINASE32 (CPK32) was identified as an interaction partner of CESA3. In this study, we provide evidence to support that CPK32 specifically phosphorylates CESA3. Overexpressing functionally defective CPK32 variant, and phospho-dead mutation of CESA3 led to deficiency in cellulose biosynthesis. Our results revealed a mechanistic understanding of the regulation of CSC stability via CPK32.

## Results

### CPK32 interacts with multiple CESAs *via* its kinase domain

The catalytic domain (amino acids residues 305 - 843) of CESA3 was used as a bait to search for putative kinases that phosphorylate CESAs. A total of 10 positive clones from Arabidopsis seedling library were identified from 100 million yeast transformants.

CPK32 was identified as a putative interaction partner of the catalytic domain of CESA3 (CESA3CD). CPK32 belongs to a multi-gene family of Ca^2+^-dependent serine/threonine protein kinases denoted CDPK/CPKs in higher plants, algae and protists (32–34). To validate the interaction between CPK32 and the CESAs, *in vitro* pull-down assays were performed. We included GST-CESA1CD in the *in vitro* pull-down assay because CESA1CD shares 77% protein sequence identity with CESA3CD. Both GST-CESA3CD and GST-CESA1CD fusion-proteins were able to pull down His-tagged full-length CPK32 (His-CPK32), which confirmed the direct interactions between CPK32 and the catalytic domain of CESAs including CESA3 and CESA1 (Fig. 1A).

**Fig. 1.**
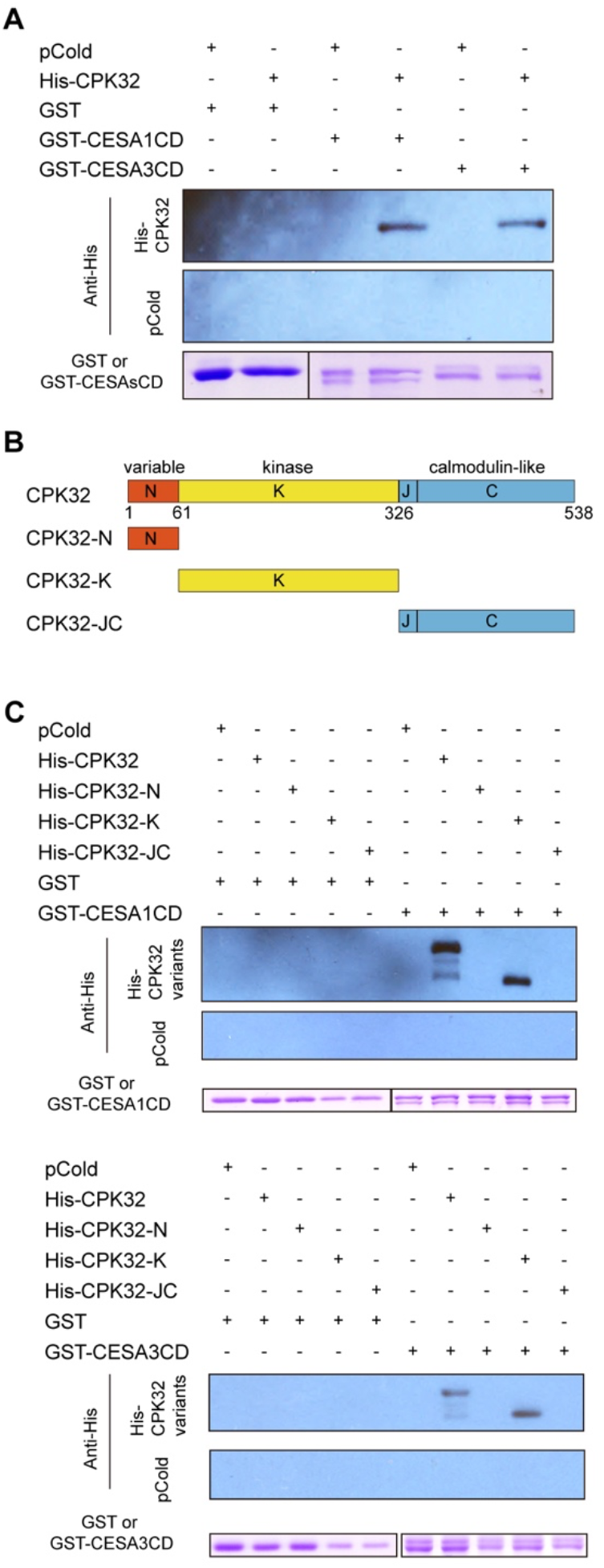
CPK32 interacts with primary CESAs *in vitro*. (A) CPK32 interacts with the catalytic domain (CD) of CESA1 and CESA3. His-CPK32 co-precipitated with both GST-CESA1CD and GST-CESA3CD in an *in vitro* pull-down assay. Empty GST beads and pCold vector without fusion genes were used as negative controls. (B) A schematic diagram displays the structure of CPK32 and its truncated fragments. N, N-terminal variable domain. K, kinase domain. J, autoinhibitory junction domain. C, calmodulin-like regulatory domain. (C) CPK32 interacts with CESAs via its kinase domain. Three CPK32 His-tagged fragments were tested for their interaction with CESAsCD *in vitro*. GST-CESA1CD and GST-CESA3CD only pulled down kinase fragment not the other fragments. Empty GST beads and pCold vector without fusion genes were used as negative controls.

CPKs share a conserved structure, consisting of an N-terminal variable domain (N), a highly conserved serine/threonine protein kinase domain (K), a short autoinhibitory junction domain (J), and a calmodulin-like regulatory domain (C) (35–38). To narrow down the interaction region of CPK32, CPK32 was divided into three fragments according to its functional domains: N, K and JC. (Fig. 1B). Both GST-CESA3CD and GST-CESA1CD were able to pull down one fragment (His-CPK32-K) but not the other two fragments (His-CPK32-N and His-CPK32-JC), indicating that the direct interaction occurs specifically between the catalytic domain of CESAs and the kinase domain of CPK32 (Fig. 1C).

### CPK32 specifically phosphorylates the catalytic domain of CESA3 *in vitro*

The direct interaction between CPK32 and CESAsCD prompted us to test whether CPK32 phosphorylates CESAsCD. Full-length CPK32 with maltose binding protein tag (MBP-CPK32) was expressed and purified in *E. coli*. The enzymatic activity was evaluated by an *in vitro* kinase assay using [γ-^32^P] ATP. As Fig. 2A shows, MBP-CPK32 has autophosphorylation activity, which is consistent with the published result (39) and served as the positive control. To eliminate the effect of trigger factor in the fusion protein, His-CESAsCD were purified and cleaved with thrombin on beads (Fig. S1). Supernatant containing cleaved products was used for the *in vitro* kinase assay. Among three different CESAsCD (CESA1CD, CESA3CD, and CESA6CD), ^32^P was only incorporated into CESA3CD (Fig. 2A), suggesting that CPK32 specifically phosphorylates the catalytic domain of CESA3 *in vitro*.

**Fig. 2.**
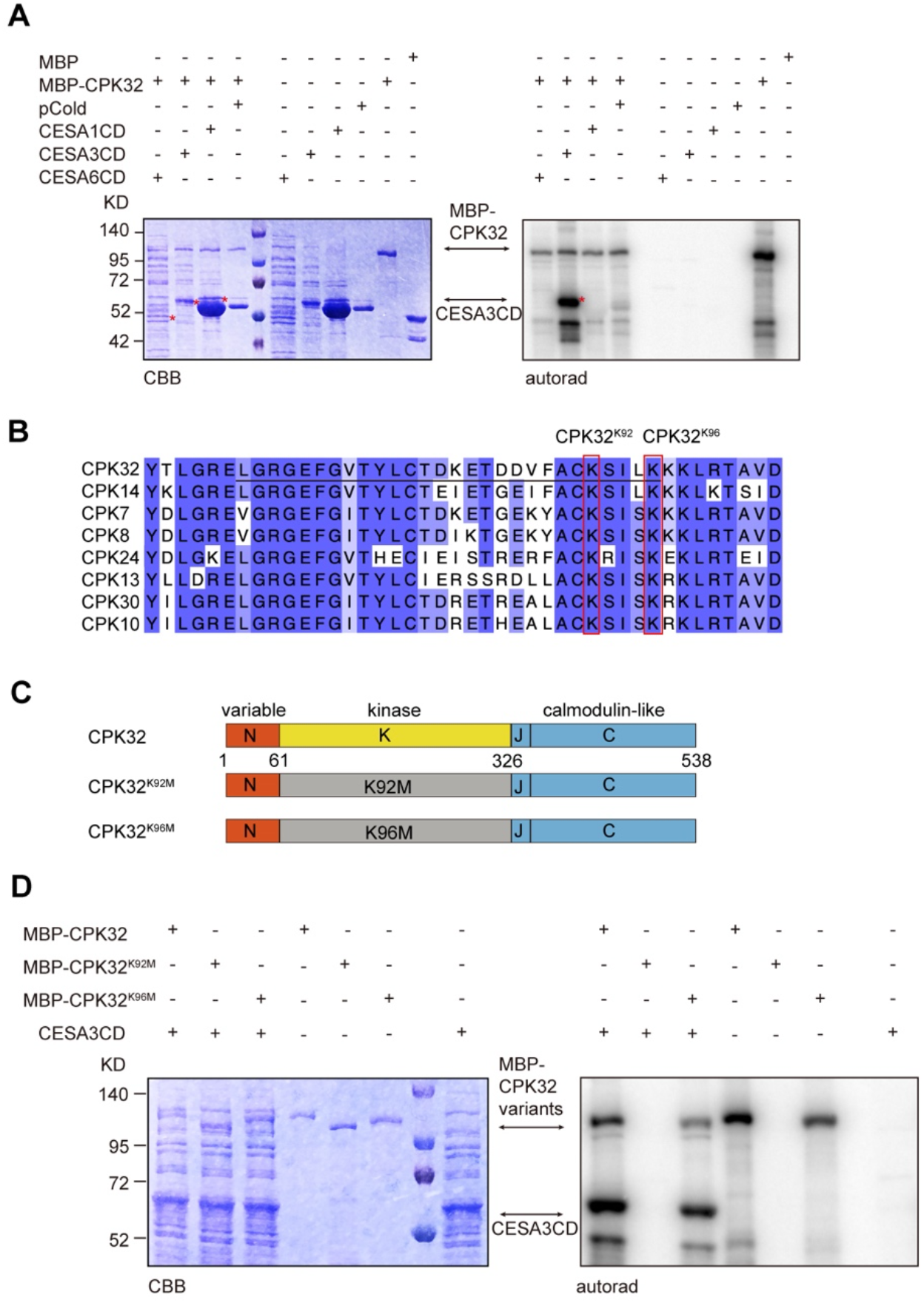
CPK32 phosphorylates CESA3 *in vitro*. (A) CPK32 specifically phosphorylates catalytic domain of CESA3. MBP-CPK32 was purified and used in *in vitro* kinase assay. His-tagged CESAsCD were purified and cleaved with thrombin before use. Left panel: Coomassie brilliant blue stained gel of recombinant proteins. Right panel: Autoradiography results of the same gel showing the signal of ^32^P. Arrows indicate the position of MBP-CPK32 and CESA3CD. Red asterisks indicate the bands of CESAsCD. This experiment was repeated three times. (B) Amino acid sequence alignment of AtCPKs in subgroup III. Putative ATP-binding region is underlined. Two conserved lysine residues are highlighted by red rectangle. (C) A schematic diagram displays the structure of CPK32 and CPK32 variants used in *in vitro* kinase assay. (D) CPK32 variants impact the kinase activity of CPK32. MBP-CPK32, MBP-CPK32^K92M^ and MBP-CPK32^K96M^ was purified and used in *in vitro* kinase assay. His-tagged CESA3CD were purified and cleaved with thrombin before use. Left panel: Coomassie brilliant blue stained gel of recombinant proteins. Right panel: Autoradiography results of the same gel showing the signal of ^32^P.

Conserved lysine residues of the ATP binding site within the kinase domain are critical for the catalytic activity of CPKs (39–41). Alignments of the amino acid sequences of 8 *Arabidopsis* CPKs in subgroup III revealed two conserved lysine residues -- Lys^92^ and Lys^96^ within the presumed ATP binding region (Fig. 2B). By mutating lysine residues to methionine, the protein kinase activity of CPK32 is expected to be disrupted (Fig. 2C). To test whether Lys-to-Met mutation affects the phosphorylation of CESA3, MBP-CPK32^K92M^ and MBP-CPK32^K96M^ were expressed and purified. MBP-CPK32^K92M^ totally abolished its kinase activity, while MBP-CPK32^K96M^ reduced both its autophosphorylation and phosphorylation of CESA3CD (Fig. 2D). Thus, Lys^92^ and Lys^96^ residues are both important for the kinase activity of CPK32 *in vitro*.

### CPK32 variants have a negative impact on the cellulose biosynthesis

A transfer DNA (T-DNA) insertion line of *CPK32* was obtained from the *Arabidopsis* Biological Resource Center, which contains a T-DNA insertion at the first exon (Fig. S2A). *cpk32-1* was verified to be a null allele as no transcripts were amplified (Fig. S2B). However, *cpk32-1* had no visible defects in overall plant growth including 4-day-old dark-grown hypocotyls, 7-day-old light-grown seedlings, and adult plants (Fig. S2C). Furthermore, the crystalline cellulose content in 4-day-old etiolated seedlings of *cpk32-1* was comparable to that of wild type (Fig. S2D). CPKs have 34 members in *Arabidopsis thaliana* that are divided into four distinct subfamilies: Group I, II, III, IV (32, 42). CPK32 belongs to the group III which has eight members: CPK7, 8, 10, 13, 14, 24, 30 and 32. It is possible that other CPKs may fulfill the function of CPK32 in *cpk32-1* null mutants. We sought to identify T-DNA mutants in Group III of CPKs. Unfortunately, we were not able to identify T-DNA null mutants for CPK8, CPK14, CPK13 and CPK24. The functional redundancy and unavailability of null mutants in Group III prevented us to generate high-order mutants to characterize the function of CPKs. Therefore, we turned to another strategy to disrupt the *in vivo* balance of CPKs and their substrates.

CPKs share a conserved activation mechanism by which the auto-inhibitory junction domain is buried within the activation site of the K domain in resting conditions, blocking its access to the substrate (43). We created three CPK32 variants: CPK32ΔC lacks J and C domains that is presumed to consistently bind to its substrate, independent of calcium regulation; CPK32ΔC^K92M^ and CPK32ΔC^K96M^ have a Lys-to-Met mutation at the ATP-binding site that were shown to disrupt the protein kinase activity (Fig. S3A). We tested whether these CPK32 variants impact the interaction with CESAs. The interaction between GST-CESA3CD and CPK32ΔC^K96M^ was comparable to that of CPK32 and CPK32ΔC (Fig. S3B). Similar results were obtained when GST-CESA1CD fusion-proteins were used in the *in vitro* pull-down assay (Fig. S3B). These results suggested that CPK32 variants do not impact their interaction with CESAs *in vitro*.

Three CPK32 variants driven by CPK32 native promoter were transformed into Col-0 or YFP-CESA6 *prc1-1*. Phenotypic assessment of the *CPK32ΔC, CPK32ΔC^K92M^* and *CPK32ΔC^K96M^* plants did not reveal severe growth defects at early developmental stages including dark-grown and light-grown seedlings (Fig. 3A). However, 7-week-old adult plants of *CPK32ΔC^K92M^* and *CPK32ΔC^K96M^* had stunted growth compared to YFP-CESA6 *prc1-1* or Col-0, while *CPK32ΔC* had comparable growth with wild type (Fig. 3A). To examine the impact of CPK32 variants on cellulose synthesis, crystalline cellulose content in 4-day-old etiolated hypocotyls of *CPK32ΔC, CPK32ΔC^K92M^* and *CPK32ΔC^K96M^* transgenic plants were measured. The results showed that *CPK32ΔC^K96M^* plants had reduced cellulose content, whereas *CPK32ΔC* and *CPK32ΔC^K92M^* were comparable to wild type (Fig. 3B). Considering MBP-CPK32^K92M^ abolished the kinase activity of CPK32 *in vitro*, it is surprising that CPK32ΔC^K92M^ did not impact cellulose production. We confirmed that mRNA level of *CPK32* in *CPK32ΔC^K92M^* and *CPK32ΔC^K96M^* transgenic lines were significantly above the native level of *CPK32* in control lines (Fig. S4A). We then examined whether similar amount of protein was produced in *CPK32ΔC^K92M^* and *CPK32ΔC^K96M^* transgenic lines. The CPK32ΔC has a C-terminal HA tag which allows us to examine the protein amount using HA antibody. As shown in Fig. S4B, CPK32ΔC^K92M^ was barely detectable in three independent transgenic lines. These results suggest that CPK32ΔC^K92M^ may affect the protein folding/stability *in vivo*. Therefore, the insignificant amount of CPK32ΔC^K92M^ might not be enough to compete against wild type copy of CPK32, which is consistent with a lack of impact in cellulose production in *CPK32ΔC^K92M^* transgenic lines.

**Fig. 3.**
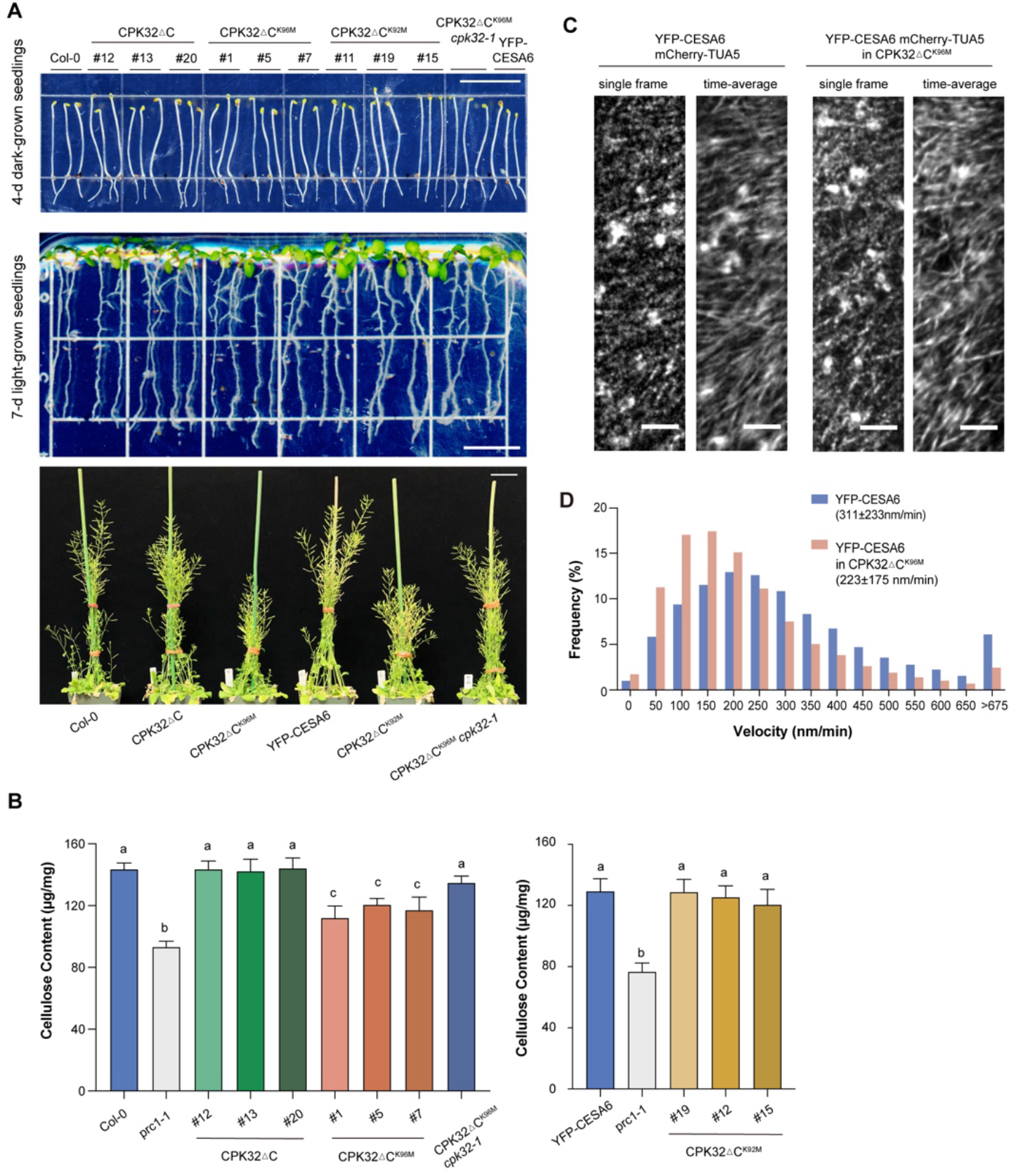
CPK32ΔC^K96M^ has a negative impact on the cellulose biosynthesis. (A) Growth phenotype of transgenic plants expressing CPK32 variants. All the transgenic plants did not exhibit visible growth phenotype at early stages including 4-day-old etiolated seedlings, 7-day-old light-grown seedlings. However, the adult plants of *CPK32ΔC^K92M^* and *CPK32ΔC^K96M^* transgenic lines had stunted growth compared to corresponding control lines and *CPK32ΔC* transgenic plants. Scale bar from top to bottom, 1 cm, 1 cm, 4 cm. (B) Crystalline cellulose content of 4-day-old etiolated seedlings of Col-0, *prc1-1*, *CPK32ΔC*, *CPK32ΔC^K96M^*, *CPK32ΔC^K96M^ cpk32-1*, *YFP-CESA6/prc1-1* and *CPK32ΔC^K92M^*. Three independent transformants of *CPK32ΔC^K96M^* plants exhibited reduced crystalline cellulose content. Error bars represent SD. Statistical analysis was performed by Two-way ANOVA Tukey’s multiple comparisons test. Lowercase letters denote statistical differences (n = 5 replicates for each line). (C) Single optical sections and time averages of 61 frames (5-min movie with 5-s intervals) of YFP-CESA6 in control lines and *CPK32ΔC^K96M^* transgenic plants. Scale bar, 5μm. (D) Histograms indicate the distribution of YFP-CESA6 particle velocities. The average CSC velocities in control and *CPK32ΔC^K96M^* transgenic line are 311 ± 233 nm/min and 223 ± 175 nm/min, respectively. n = 19,481 particles from 11 regions of interest (ROIs) in 6 individual seedlings for control line (YFP-CESA6 mCherry-TUA5 in *prc1-1)*. n = 28,234 particles from 14 ROIs in 10 individual seedlings for *CPK32ΔC^K96M^* transgenic line (YFP-CESA6 mCherry-TUA5 in *CPK32ΔC^K96M^ prc1-1)*.

To examine whether the CPK32 variants influence CSC dynamics, *CPK32ΔC^K96M^* plant was crossed with an *Arabidopsis* transgenic line expressing both YFP-CESA6 and mCherry-TUA5. Quantification of the CSC velocity revealed that the CSC motility was decreased in *CPK32ΔC^K96M^* compared to that in control seedlings. The CSCs had an average velocity of 311 ± 233 nm/min (n = 19,481) in control cells. However, with the expression of CPK32ΔC^K96M^, the average velocity of CSCs had a 28.3% reduction to 223 ± 175 nm/min (n = 28,234) (Fig. 3C and D). The reduction in CSC motility in *CPK32ΔC^K96M^* plants is consistent with the reduced cellulose content.

### CPK32 modulates the dynamics of CSC via phosphorylation of CESA3

Phosphoproteomic studies have identified multiple phosphorylation sites on CESA3, including S671 in the catalytic domain of CESA3 (44). To test whether CPK32 phosphorylates CESA3 on specific residue in the catalytic domain, we performed *in vitro* kinase assay with the same experimental setting as Fig. 2A, except for supplying regular ATP. The phosphorylated CESA3CD protein was separated by SDS-PAGE and the protein bands between 72 KD and 52 KD were cut for in-gel trypsin digestion prior to liquid chromatography-tandem mass spectrometry (LC-MS/MS). Contrary to PhosPhat 4.0 phosphorylation database, T672 was identified to be phosphorylated in the assay group (CPK32 + CESA3CD + ATP) but not in the negative control group (CPK32 + CESA3CD) (Fig. S5). Because S671 and T672 are next to each other, we decided to mutate both S671 and S672 to alanine. We generated binary vector containing GFP tagged CESA3^S671A T672A^ under the control of *CESA3* promoter and transformed it into heterozygous *cesa3-1* mutant, as the *cesa3-1* knock-out mutant is gametophytic lethal (20). As a control, the GFP tagged wild type copy of CESA3 was transformed into segregating *cesa3-1* mutant background. Transgenic plants expressing GFP-CESA3 or GFP-CESA3^S671A T672A^ were identified by hygromycin resistance. PCR was performed to identify *cesa3-1* homozygous lines in the T2 generation.

Similar to *CPK32ΔC^K96M^* transgenic plants, *GFP-CESA3^S671A T672A^* transgenic lines in *cesa3-1* null background were phenotypically indistinguishable from *GFP-CESA3* control lines during the seedling growth (Fig. 4 A and B). The crystalline cellulose content in 10-day-old light-grown seedlings of *GFP-CESA3* and *GFP-CESA3^S671A T672A^* were measured to assess the cellulose synthesis. The result showed that *GFP-CESA3^S671A T672A^* plants had reduced cellulose content compared to the control *GFP-CESA3* lines, whereas *GFP-CESA3* was comparable to wild type (Fig. 4C). Consistently, quantification of the CSC velocity indicated that the CSC motility was decreased in *GFP-CESA3^S671A T672A^* plants compared to that in control line. The CSCs had an average velocity of 187 ± 90 nm/min (n = 1,782) in control cells. However, with the GFP-CESA3^S671A T672A^, the average velocity of CSCs had a 27.8% reduction (p<0.0001) to 135 ± 85 nm/min (n = 2,532) (Fig. 4 D and E). These results indicated that CPK32 regulates the CSC dynamics and cellulose synthesis by the phosphorylation of CESA3 on S671 and T672 residues.

**Fig. 4.**
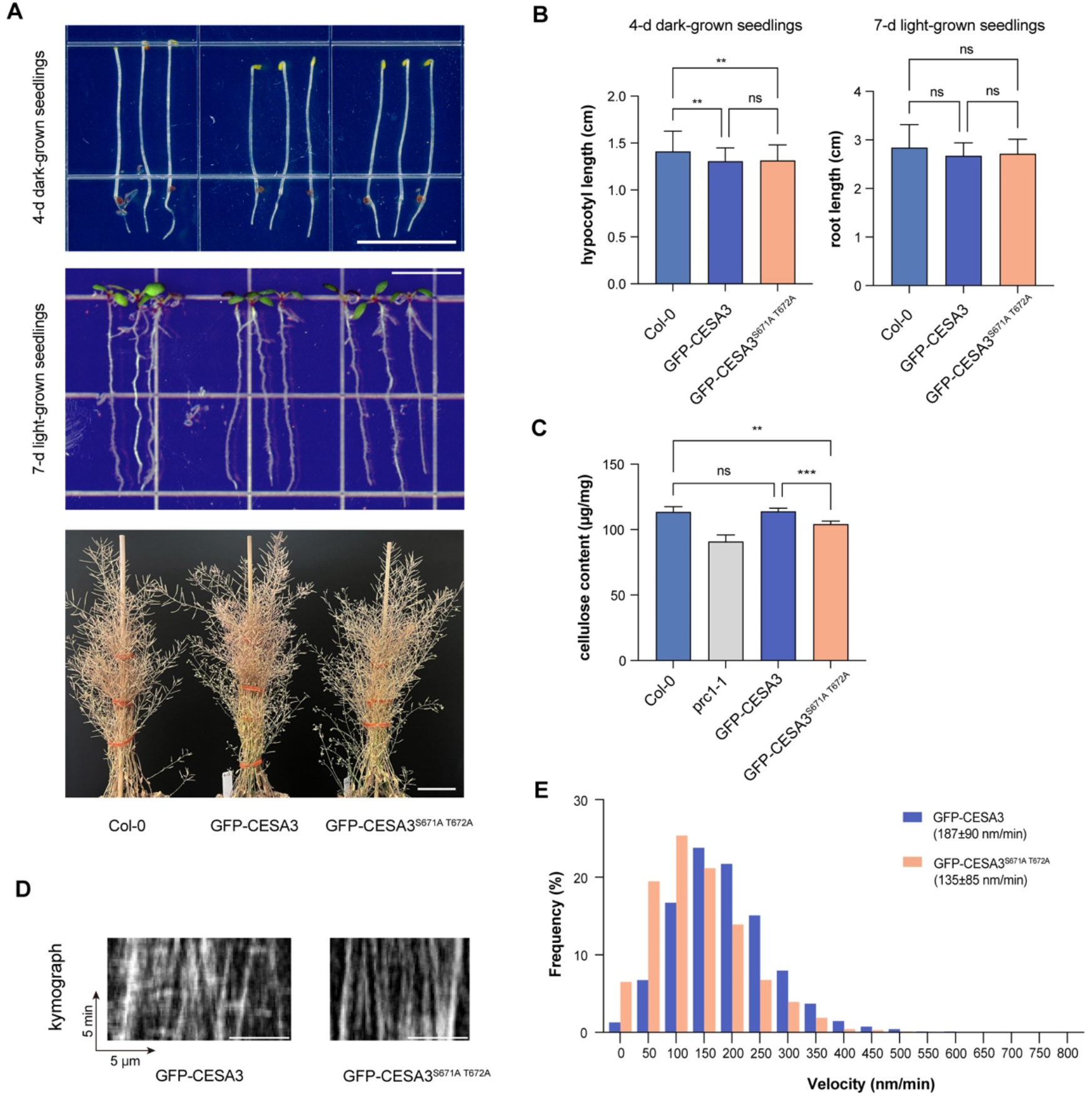
*GFP-CESA3^S671A T672A^* transgenic lines showed impaired cellulose biosynthesis. (A) Growth phenotype of transgenic plants expressing GFP-CESA3 and GFP-CESA3^S671A T672A^. *GFP-CESA3* and *GFP-CESA3^S671A T672A^* complemented *cesa3-1* and showed comparable growth for 4-d etiolated seedlings. 7-d light-grown seedlings and adult plants were indistinguishable between Col-0 and GFP-CESA3^S671A T672A^. Scale bar from top to bottom, 1 cm,1 cm and 4 cm. (B) Measurement of the hypocotyl length of 4-d etiolated seedlings and the root length of 7-d light-grown seedling of Col-0, GFP-CESA3 and GFP-CESA3^S671A T672A^. Error bars are SD. **P < 0.01. (C) Crystalline cellulose content of 10-d light-grown seedlings of Col-0, GFP-CESA3 and GFP-CESA3^S671A T672A^. GFP-CESA3^S671A T672A^ showed reduced crystalline cellulose content. Error bars represent SD. Statistical analysis was performed by Two-way ANOVA Tukey’s multiple comparisons test. **P < 0.01, ***P<0.001 (n = 6 replicates for each line). (D) Kymographs derived from 5-min movies with 5-s intervals. (E) Histograms indicate the distribution of GFP-CESA3 or GFP-CESA3^S671A T672A^ particle velocities. The average velocity of GFP-CESA3 is 187 ± 90 nm/min (n = 1,782 particles from 17 regions of interest (ROIs) in 11 individual seedlings) and that of GFP-CESA3^S671A T672A^ is 135 ± 85 nm/min (n = 2,532 particles from 18 regions of interest (ROIs) in 8 individual seedlings).

### CPK32 regulates the stability of CSCs

CSC phosphorylation was shown to regulate motility and bidirectional movement of CSC in primary cell walls (26–28). In addition, it was long proposed that phosphorylation may regulate the formation of CSCs, protein stability, and/or activation of CSCs at the plasma membrane (19, 45). We examined whether the phosphorylation impact the assembly of CSCs by performing blue-native polyacrylamide gel electrophoresis analyses of accumulated microsomal fraction of seedlings expressing CPK32ΔC^K96M^ and corresponding control lines. There was no detectable difference in band intensities in *CPK32ΔC^K96M^* plants compared with that of wild type, indicating that CPK32ΔC^K96M^ does not impact the assembly of CSCs (Fig. S6).

We then tested whether phosphorylation has a role in regulating the stability of CSCs. Previous study showed that the decay of CESA1, 3 and 6 was equivalent over time (46). To assess protein stability *in vivo*, we analyzed the decay of YFP-CESA6 in the seedlings overexpressing CPK32ΔC^K96M^ (CPK32ΔC^K96M^ YFP-CESA6 mCherry-TUA5/*prc1-1*) and corresponding control line (YFP-CESA6 mCherry-TUA5/*prc1-1*). CESAs in light-grown seedlings had a half-life of around 36 h and around 20% of CESA proteins remained after 48 h of 1 mM cycloheximide treatment (46). We used same experimental settings and analyzed CESA decay at four time points: 0 h, 24 h, 36 h, and 48 h. 44.0% of YFP-CESA proteins remained after 36 h of cycloheximide treatment in the seedlings overexpressing CPK32ΔC^K96M^, compared to 75.4% in control line (Fig. 5). These results indicated that deregulation of CPKs impacts the stability of CSCs, possibly due to deficiency in the phosphorylation of CESA3.

**Fig. 5.**
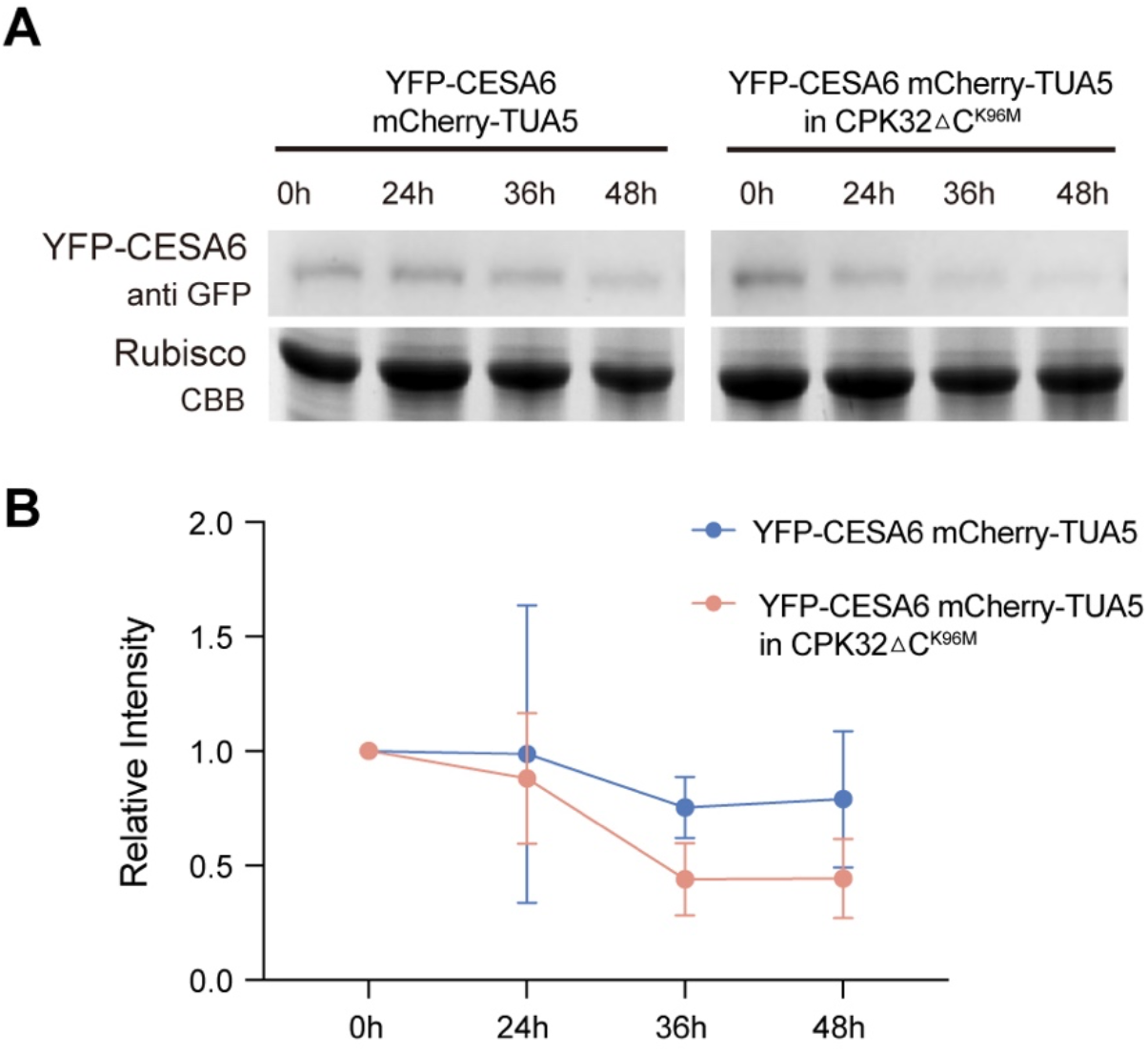
Decay of YFP-CESA6 was accelerated in *CPK32ΔC^K96M^* line. (A) Time-dependent decay of YFP-CESA6 in control (YFP-CESA6 mCherry-TUA5) and *CPK32ΔC^K96M^* transgenic plants. Upper panel shows the immunoblot and bottom panel shows the loading control. (B) The band intensities were quantified and normalized to T=0. The mean of adjusted band intensity of YFP-CESA6 from control and *CPK32ΔC^K96M^* plants were obtained at each time points (n=5). Bars indicate SD.

## Discussion

Despite genetic and cell biological evidence showing the importance of CESAs phosphorylation, we have limited knowledge of the corresponding kinase and regulatory mechanism. In this study, CPK32 was identified as the interaction partner of CESA3. The biological functions of CPK32 have been reported in many biological processes, such as pollen tube growth, shoot and root development and ABA-induced transcriptional regulation (39, 47–49). Our study adds a novel function of CPK32 that regulates the stability of CSCs via phosphorylation of a specific CESA isoform CESA3. CPK32 directly interacts with the catalytic domains of multiple CESAs including CESA1 and CESA3. However, among three substrates including CESA1, CESA3, and CESA6, CPK32 specifically phosphorylates CESA3 on T672 residue *in vitro*. CESA3^T672^ is located at variable region 2 within the catalytic domain, also known as class-specific region (CSR). CSR shares limited sequence conservation among paralogous CESAs (23, 44, 50, 51). The cryo electron microscopy (cryo-EM) structure of PttCESA8 indicates that disordered CSR domain may affect interactions with other binding partners (52). Many phosphorylation sites localize to CSR domain. For example, both Ser^686^ and Ser^688^ of CESA1 reside in CSR domain and are important for anisotropic cell expansion and cellulose synthesis (27). Ser^686^ and Ser^688^ of CESA1 is next to CESA3^T672^ within the CSR. However, CPK32 did not phosphorylate CESA1 *in vitro*.Given that phosphoproteomic studies revealed various phosphorylation sites among CESAs, it is likely that the regulation of CESAs involves different kinases.

Functional redundancy of CPKs is common due to gene duplication within the CPK subgroup (39, 47, 49). For instance, it has been reported that CPKs function redundantly in nitrate-signaling. CPK10, CPK30, and CPK32 phosphorylate the transcription factor, NIN-LIKE PROTEIN7 (NLP7), *in vitro*. The *cpk10*, *cpk30*, and *cpk32* single mutants barely affected the expression of nitrate-responsive genes. The nitrate-responsive genes were reduced in *cpk* double mutants and further exaggerated in the conditional triple mutants (47). We proposed that multiple CPKs might modulate cellulose synthesis. Consistent with our hypothesis, *cpk32-1* null mutant had no visible defect in cellulose production. *CPK32ΔC^K96M^* transgenic plants in the wild-type background had reduced CSC motility and reduced cellulose content, presumably due to a dominant negative effect on sequestering CESA3 and impaired phosphorylation of CESA3. This is consistent with the observation that CPK32 variants do not impact their interaction with CESA3 *in vitro*. Interestingly, we observed that when CPK32ΔC^K96M^ was introduced to *cpk32-1* null mutant, CPK32ΔC^K96M^ was not able to impact cellulose production (Fig. 3 A and B). It is likely CPK32 may homodimerize itself or heterodimerize with other CPKs. Future research will be directed towards a mechanistic understanding of sub-functionalization of CPKs.

By mutating both CESA3^S671^ and CESA3^T672^ to alanine, it is predicted that CESA3 would not be regulated by CPKs. Indeed, *GFP-CESA3^S671A T672A^* transgenic lines phenocopied aspects of the *CPK32ΔC^K96M^* lines such as reduced cellulose content and impaired CSC motility. However, *GFP-CESA3^S671A T672A^* transgenic lines did not show stunted adult plants. It is likely that CPK32ΔC^K96M^ would impact the balance of other CPKs and additional substrates. One possibility is CPKs that modulate shoot development growth (47). The similar phenotype of *GFP-CESA3^S671A T672A^* and *CPK32ΔC^K96M^* lines indicated that CPK32 facilitates the specific phosphorylation of CESA3^T672^ and thus influences the CSCs motility and cellulose biosynthesis.

Phosphorylation plays an important role in the regulation of CSCs velocity, bidirectional movement, and anisotropic cell expansion. Disruption a balance of phosphorylation and dephosphorylation by introducing phospho-mimic glutamine (E) or phospho-dead alanine (A) revealed the regulation of CESA via its post-translational modification. For example, residues in the hypervariable region of CESA1^T165A^ and CESA3^S211A^ are important for maintaining anisotropy and bi-directional microtubule-based motility (26, 27). However, the molecular mechanisms of CESA phosphorylation are unknown. Our study revealed that phosphorylation regulates the stability of CSCs.

The degradation of CESA proteins in CPK32ΔC^K96M^ was accelerated, indicating that phosphorylation and dephosphorylation of CESA3 regulates the stability of CSCs. In primary cell walls, CSCs are unusually stable with half-life over 48 hours (46). Cycloheximide removed majority of CSCs from the plasma membrane, but Golgi-localized CSCs were retained after 50 hours of 1 mM cycloheximide treatment (46). CSCs have been shown to cycle between Golgi apparatus, small CESA compartment or microtubule-associated cellulose synthase compartments (SmaCCs/MASCs), and plasma membrane (53). The number of CSCs in these compartments is expected to undergo tight regulation. Under abiotic stress, CSCs are internalized via clathrin-mediated endocytosis (53). Internalized CESAs can be sorted to SmaCCs/MASCs for endocytic recycling, to the Golgi for reassembly, or to the vacuole for degradation. It is unclear which routes of CSCs trafficking CPK32ΔC^K96M^ affects. Phosphorylation could result in conformational changes that induce the disengagement of CSCs from the cortical microtubules, thereby acting as a cue for clathrin-mediated endocytosis. It remains to be empirically tested whether phosphorylation influences the endocytosis of CESAs. Phosphorylation of two non-conserved residues of CESA7 has been associated with proteosome-dependent degradation of CESA7 (19). Further studies are required to determine the impact of phosphorylation in the trafficking route and degradation process of CESAs.

## Materials and Methods

### Yeast two-hybrid assay

The catalytic domain of CESA3 (CESA3CD) was cloned into the bait vector pAS2 containing the *GAL4* DNA binding domain using primers shown in Table S1. pAS2-CESA3CD was introduced into Y190 by electroporation, resulting in the Y190 pAS2-CESA3CD clone. After testing for no auto-activation when combined with pACT vector containing the GAL4 DNA activation domain, Y190 pAS2-CESA3CD was transformed with ~10 μg of the Arabidopsis thaliana seedling cDNA library. Yeast transformants were plated on selection medium lacking tryptophan, leucine, histidine and supplemented with 100 mM 3-aminotriazole. A filter assay was performed to test for the ß-galactosidase activity on all transformants. A total of 10 positive colonies were recovered and subjected for sequencing using the T7 primer.

### Plant Materials and Growth Conditions

T-DNA insertion lines of *cpk32-1* (GABI_824E02) and *cesa3-1* (SALK_014134) were obtained from the Arabidopsis Biological Resource Center. Primers used for genotyping are listed in Table S1.

To create *pCPK32::CPK32-YFP* binary construct, the HA tag was removed from pEarleyGate 301 (TAIR) and replaced by an YFP tag, result in a modified version of pEarleyGate 301. DNA fragment containing a 2-kb CPK32 promoter and the genomic sequence of CPK32 were PCR-amplified from the genomic DNA, cloned into pDONR/Zeo vector (Thermofisher). After sequence verification, the pDONR/Zeo *pCPK32::CPK32* was introduced into the modified pEarleyGate 301 via Gateway cloning. To create *pCPK32::CPK32ΔC-HA* binary construct, a DNA fragment containing a 2-kb CPK32 promoter and the genomic sequence of CPK32 (without junction and calmodulin domain) were PCR-amplified from the genomic DNA using the primers listed in Table S1, cloned into pDONR/Zeo vector, sequenced, and introduced into the original pEarleyGate 301 via Gateway cloning. To generate *pCPK32::CPK32ΔC^K92M^-HA and pCPK32::CPK32ΔC^K96M^-HA* binary constructs, lysine residues were mutated to methionine via site-directed mutagenesis on the *pCPK32::CPK32ΔC-HA* template. To generate *pCESA3::GFP-CESA3* binary construct, the coding sequence of CESA3 was PCR-amplified from the cDNA library, cloned into pCR8/GW/TOP vector. After sequence verification, CESA3 was introduced into the modified pH7WGF2 vector whose original promoter was replaces by a 1831bp CESA3 promoter. To generate *pCESA3::GFP-CESA3^S671A T672A^* binary construct, serine and threonine residues were mutated to alanine via side-directed mutagenesis on the *pCESA3::GFP-CESA3* template. All the binary constructs were introduced into corresponding background by *Agrobacterium tumefaciens-mediated* transformation using the floral dipping method. *pCPK32::CPK32-YFP* was introduced into *cpk32-1. pCPK32::CPK32ΔC-HA* and *pCPK32::CPK32ΔC^K96M^-HA* were introduced into Col-0. *pCPK32::CPK32ΔC^K92M^-HA* was introduced into YFP-CESA6/*prc1-1*. *pCESA32::GFP-CESA3* and *pCESA3::GFP-CESA3^S671A T672A^* were introduced into heterozygous *cesa3-1*. All the primers used are listed in Table S1.

All seeds were surface sterilized with 30% (vol/vol) bleach for 15 min, thoroughly washed with sterile double-distilled H_2_O, and stored at 4 °C for a minimum of 3 days. Etiolated seedlings were grown on vertical half-strength Murashige and Skoog (MS) plates [1/2 × MS salts, 0.05% mono-hydrate 2-(N-Morpholino) ethanesulfonic acid, 0.8% agar, pH 5.7] at 21 °C in the dark. Light-grown seedlings were grown on vertical half-strength MS plates with 1% sucrose at 21 °C on a 16-h light/8-h dark cycle.

### Protein Purification

The coding sequences of the genes were cloned into pCold-TF vector (TaKaRa) which contains a His tag, or pGEX-KG vector in frame with a GST tag, or pMAL-C2 vector (NEB) in frame with an MBP tag. Fusion genes were expressed in BL21-CodonPlus (DE3)-RIPL *Escherichia coli* (*E. coli*). Fusion genes in pCold-TF were induced with 1 mM isopropyl β-D-1-thiogalactopyranoside (IPTG) at 18 °C for 16 h after a 30 min 15°C cold shock. Fusion genes in pGEX-KG or pMAL-C2 were directly induced with 1 mM isopropyl β-D-1-thiogalactopyranoside (IPTG) at 18 °C for 16 h. Protein purification was performed as described previously (54). All the primers used are listed in Table S1.

### *In vitro* Pull-down Assay and Western Blot Analysis

Resin-bound GST-fusion proteins were washed twice with interaction buffer (20 mM HEPES, pH 7.4, 1 mM EDTA, 5 mM MgCl_2_, 1 mM dithiothreitol, and 0.1% Triton X-100) for equilibration. Aliquots of approximately 2 μg of equilibrated resin-bound GST-fusion proteins were incubated with approximately 2 μg of soluble His-tagged proteins in a total volume of 500 μl of interaction buffer for 2h at 4°C on a rocker. The resin was then washed eight times with interaction buffer, resuspended in 2x Laemmli protein sample buffer (BIORAD), boiled, and subjected to SDS-PAGE and western blotting. On western blots, His-tagged proteins were detected on film by chemiluminescence using a horseradish peroxidase-conjugated His antibody and SuperSignal West Femto substrate (ThermoFisher).

### *In vitro* Protein Kinase Assay

To cleave the His tag and trigger factor, the resin-bound recombinant proteins were incubated with 5 μl thrombin (1U/μl) at room temperature for 2h. The supernatants containing cleaved proteins were used in protein kinase assay. Because the MBP tag did not affect the enzymatic activity of CPK32, MBP tag was remained after protein purification. The kinase assay was performed at 25°C for 5 min according to the reference (39). Specifically, 0.3 μg of the purified MBP-CPK32 or MBP-CPK32 variants was incubated with 0.6 μg of CESAsCD in 25 μl of kinase buffer (25 mM Tris-HCl, pH 7.5, 10 mM MgCl2, 10 mM ATP) containing 2 μCi of [γ-^32^P]ATP. CPK32 autophosphorylation was carried out in 25 μl of kinase buffer in the presence of 1 μg of MBP tagged CPK32 or CPK32 variants at 25°C for 20 min.

The products after reaction were separated by SDS-PAGE gel and stained with Commassie brilliant blue. The SDS-PAGE gel was then dried by BIO-RAD Gel Dryer (model 583), exposed radioactivity to storage phosphor screen (BIO-RAD sample exposure cassette and phosphor screen) overnight and visualized by GE Healthcare Typhoon 9410.

### In-gel digestion

SDS-PAGE gel bands were reduced with 10mM DTT for 30 min at 60°C, alkylated with 20mM iodoacetamide for 45min at room temperature in the dark and digested overnight with 0.2μg of either trypsin (37°C), chymotrypsin (20°C) or AspN (37°C). All proteases were Pierce MS Grade (ThermoFisher). Peptides were extracted twice with 5% formic acid, 60% acetonitrile and dried under vacuum.

### Liquid chromatography-tandem mass spectrometry (LC-MS/MS)

LC-MS/MS was conducted using a nano LC (Dionex Ultimate 3000 RLSCnano System, ThermoFisher) interfaced with an Eclipse Tribrid mass spectrometer (ThermoFisher). Each sample 1/20 of digests) was loaded onto a fused silica trap column (Acclaim PepMap 100, 75 μm × 2 cm, ThermoFisher). After washing for 5 min at 5 μl/min with 0.1% TFA, the trap column was brought in-line with an analytical column (Nanoease MZ peptide BEH C18, 130A, 1.7μm, 75μm×250mm, Waters) for LC-MS/MS. Peptides were fractionated at 300 nL/min using a segmented linear gradient of 4-15% B in 5 min (where A: 0.2% formic acid, and B: 0.16% formic acid, 80% acetonitrile), 15-50% B in 50 min, and 50-90% B in 15min. Solution B is then returned to 4% for 5 minutes before the next run.

The scan sequence began with an MS1 spectrum (Orbitrap analysis, resolution 120,000, scan range from M/Z 275–1500, automatic gain control target 1E6, maximum injection time 100 ms). The top 5 (3 sec) duty cycle scheme was used to determine the number of MS/MS scans performed for each cycle. Precursor ions of charges 2-7 were selected for MS/MS and a dynamic exclusion of 60sec was used to avoid repeat sampling. Precursor ions were isolated in the quadrupole with an isolation window of 1.2 m/z, automatic gain control target 1E5, and fragmented with higher-energy collisional dissociation with a normalized collision energy of 30%. Fragments were scanned in Orbitrap with resolution of 15,000. MS/MS scan ranges were determined by the charge state of the parent ion but a lower limit was set to 110 m/z.

### Database Analysis

Nano LC-MSMS Raw file was analyzed by Proteome discoverer (v 2.4.1.15) using Sequest HT search engine against E.coli K12 database (NCBI) as well as custom supplied sequences and a database composed of common lab contaminants. Digestion protease was set as trypsin/chymo/Asp-N full digestion. Precursor mass tolerance was set at 10 ppm and fragment mass tolerance were set at 0.02 Dalton. Carboxyiodomethyl on cysteine was set as static modification, acetylation, methionine loss and acetylation plus methionine loss were set as dynamic modifications at the protein terminus. Phosphorylation on serine, threonine and tyrosine as well as oxidation on methionine was set as dynamic modification. Percolator was used to validate results with strict target false discovery rate set to 0.01 and relaxed target rate set to 0.05 for both peptides and proteins. Phospho-RS was used to calculate phosphorylation sites possibilities. Precursor ion intensity was used to represent peptide and protein abundance. Peptides were grouped into protein groups using strict parsimony principle.

### RNA Isolation and Quantitative RT-PCR Analysis

RNA was purified from seedlings Col-0 and *cpk32-1* using the Direct-zol RNA MiniPrep Kit (Zymo Research). The reverse-transcriptase cDNA synthesis reaction was carried out by the RevertAid Reverse Transcriptase (ThermoFisher).

For qPCR analysis, cDNA from each genotype was subjected to PCR using primer pairs (Table S1) that were specific to the coding sequence of *CPK32* and *ACTIN2* for 24 cycles and analyzed by gel electrophoresis. For Real-Time quantitave PCR (RT-qPCR) analysis, Luna^®^ Universal qPCR Master Mix (NEB) was used for qPCR on an ABI StepOne Plus real-time system (Life Technologies). RT-qPCR was performed in triplicate and data were collected by ABI STEPONETM software version 2.1. The relative expression of each gene was normalized to *ACTIN2* gene (AT3G18780) expression.

### Cellulose Content Measurement

4-day-old etiolated seedlings from half-strength MS plates without sucrose or sample described previously were collected. The crystalline cellulose was measured using the Updegraff method (55). Data were collected from at least five technical replicates for each genotype.

### Microsomal Fraction Isolation and Native Protein Electrophoresis

7-d light-grown seedlings are collected, grounded and resuspended with lysis buffer (2mM EGTA, 2mM EDTA, 100mM MOPs pH 7.0, Roche protease and phosphatase Inhibitor cocktail tablets). The crude extraction was centrifuged at 5,000 g for 10 min. The supernatant was further centrifuged at 4 °C, 100,000 g for 1 hour. The supernatant was collected and marked as ‘supernatant’ in Fig. S6. The pellet (crude microsomal pellet) was collected and resuspended in 150 μl resuspension buffer (lysis buffer and 2% (v/v) of Triton X-100). After incubation on ice for 30 min, the mixture was centrifuged at 4 °C, 100,000 g for 30 min. The supernatant was marked as ‘pellet’ in Fig. S6 and subjected for BN-PAGE electrophoresis using the NativePAGE Novex 3–12% Bis-Tris Protein Gels (Invitrogen), following the protocol provided by Invitrogen. For immunoblot, the gel was treated with denaturing buffer (3.3% (m/v) SDS, 65 mM Tris-HCl, pH6.8) and subjected to western blot analysis.

### Live-cell Imaging and Analysis

Images were obtained from epidermal cells of 2.5-d-old, etiolated seedlings about 0.5 - 2 mm below the apical hook. Imaging was performed on a Yokogawa CSUX1 spinning-disk system as previously described (56). For colocalization analyses, CESA particle dynamics, images were collected as previously described (57). Image analysis was performed using Metamorph, Imaris, and ImageJ software.

### Protein Degradation Assessment

The assay was adapted from previous research (46). 7-day light-grown seedlings were treated in liquid half-strength MS medium in the presence of 1% sucrose and 1mM CHX for 0, 24, 36, 48 hours. The medium was refreshed every 24 h to avoid the degradation of CHX. The seedlings were grounded and resuspended in lysis buffer (50 mM Tris pH7.5, 150 mM NaCl, 0.5% Triton X-100) and incubated on ice for 30min. The supernatant after spin (13000 rpm 30min at 4 °C) was collected and assessed for protein amount by Bradford Assay (58). 50 μg protein of samples were loaded to assess the YFP-CESA6 by immunoblot using anti-GFP (Invitrogen, Cat# A-11122). 5 μg protein of samples were loaded to adjust the measurement. The band intensity of YFP-CESA6 and Coomassie brilliant blue staining Rubisco were measured by ImageJ’s gel analysis.

## ACKNOWLEDGMENTS

We gratefully acknowledge support during the initiation of this research from Christopher R. Somerville and the Energy Biosciences Institute. W.D., X.X., L.L. was supported by the Center for Lignocellulose Structure and Formation, an Energy Frontier Research Center funded by the US Department of Energy, Office of Science, Basic Energy Sciences, under Award DE-SC0001090. S.L. was supported by the startup funds from Department of Biochemistry and Molecular Biology, Pennsylvania State University. Y.G. was supported by the National Science Foundation (NSF) #1951007.

## Supplementary Figures

**Supplementary Fig. S1.**
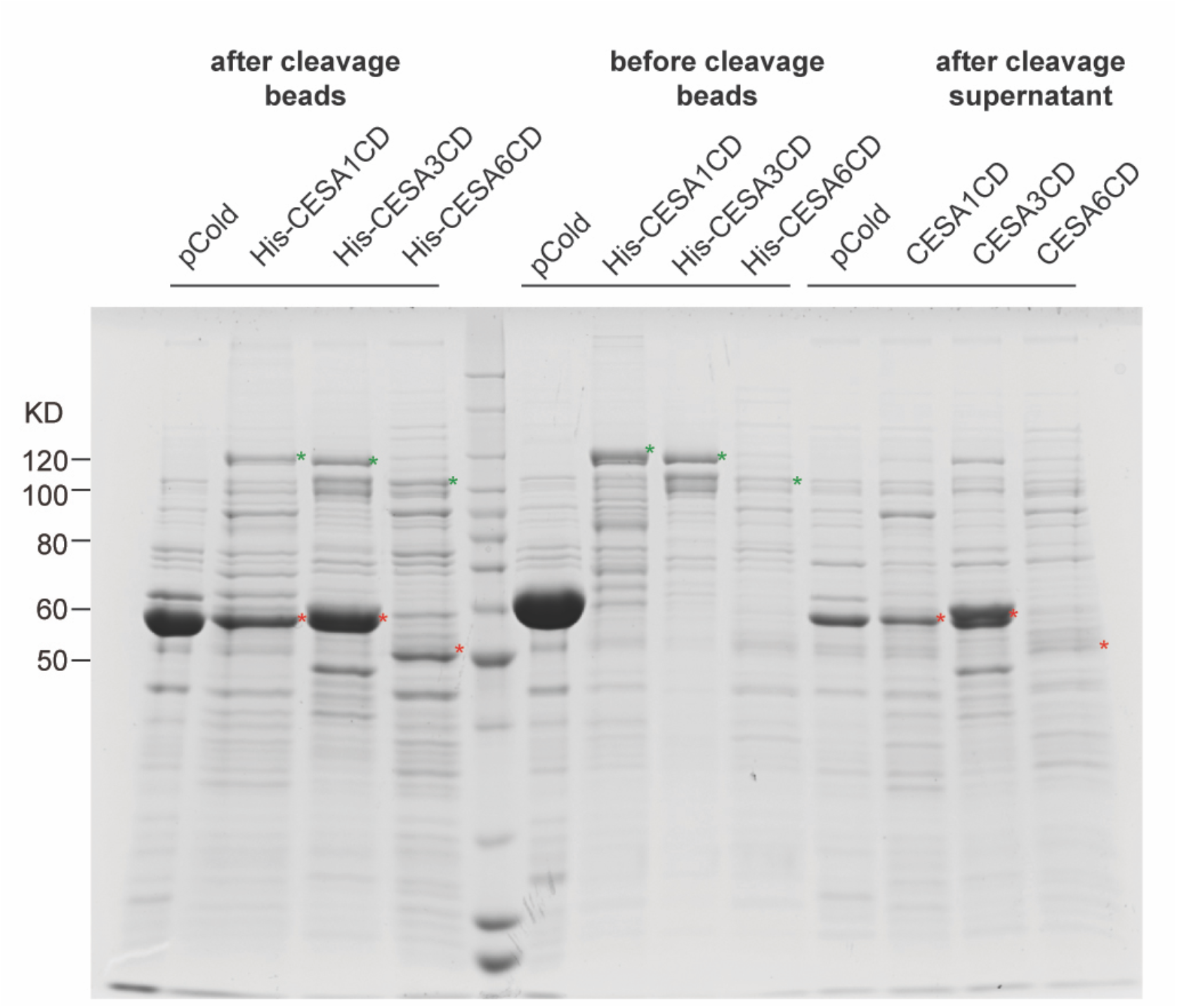
Purification and cleavage of His-CESAsCD for *in vitro* kinase assay. The Coomassie brilliant blue stained gel shows purified His-CESAsCD and their products after thrombin cleavage. Resin-bound recombinant proteins were incubated with 5 μl thrombin (1U/μl) at room temperature for 2h. Green asterisks indicate the expected size of His-CESAsCD. Red asterisks indicate the expected size of CESAsCD after cleavage. Supernatant containing cleaved CESAsCD was used for *in vitro* kinase assay.

**Supplementary Fig. S2.**
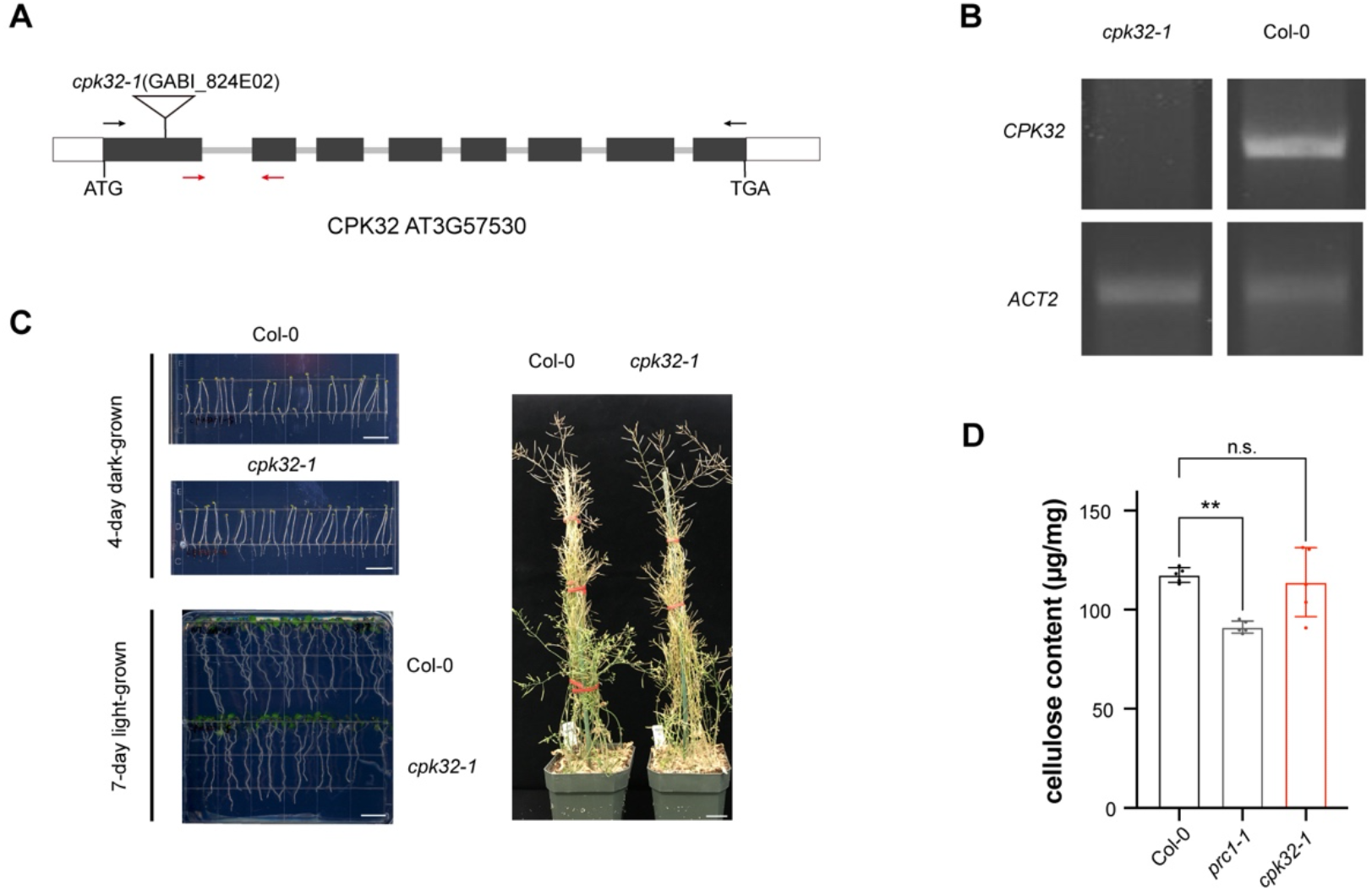
*cpk32* single mutant display no visible growth phenotype. (A) The gene model of *CPK32* shows the position of T-DNA insertion in *cpk32-1* mutant. Black arrows indicate the primer pair for qPCR. Red arrows indicate the primer pair for RT-qPCR. (B) Quantitative-PCR analysis of *CPK32* expression from wild type and *cpk32-1* mutant. Primers are indicated by black arrows in Fig. S1A. *ACTIN2 (ACT2)*was used as the reference gene. (C) Phenotype of wild type (Col-0) and *cpk32-1* of 4-d etiolated seedlings, 7-d light-grown seedlings, and adult plants. Scale bar, 1 cm in left panel and 2 cm in the right panel. (D) Crystalline cellulose content of 4-d etiolated seedlings of wild type (Col-0), *prc1-1*,and *cpk32-1*. Error bars are SD. Statistical analysis was performed by one-way ANOVA Tukey test (n = 5 replicates for each genotype).

**Supplementary Fig. S3.**
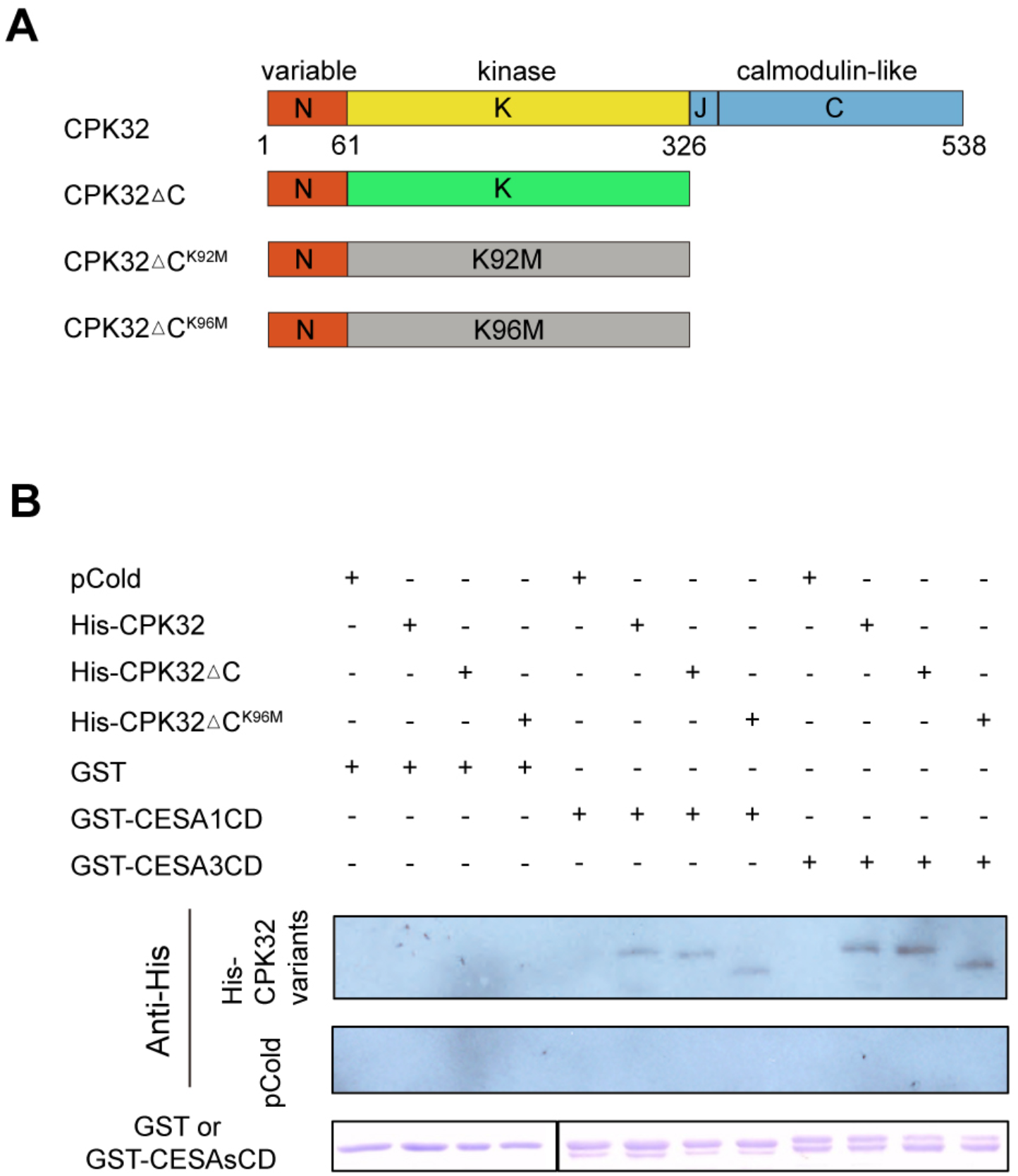
CPK32 variants remain the *in vitro* interaction with CESAsCD. (A) A schematic diagram of the CPK32 variants. CPK32ΔC was generated by truncating the auto-inhibitory junction domain (J) and a C-terminal calmodulin-like domain (C). A Lys (K) to Met (M) mutation was created to generate CPK32ΔC^K92M^ and CPK32ΔC^K96M^ variants. (B) CPK32ΔC and CPK32ΔC^K96M^ interact with CESAs. Both GST-CESA1CD and GST-CESA3CD were able to pull down His tagged CPK32ΔC and CPK32ΔC^K96M^. His-CPK32 served as positive control. GST alone and pCold vector without fusion genes were used as negative controls.

**Supplementary Fig. S4.**
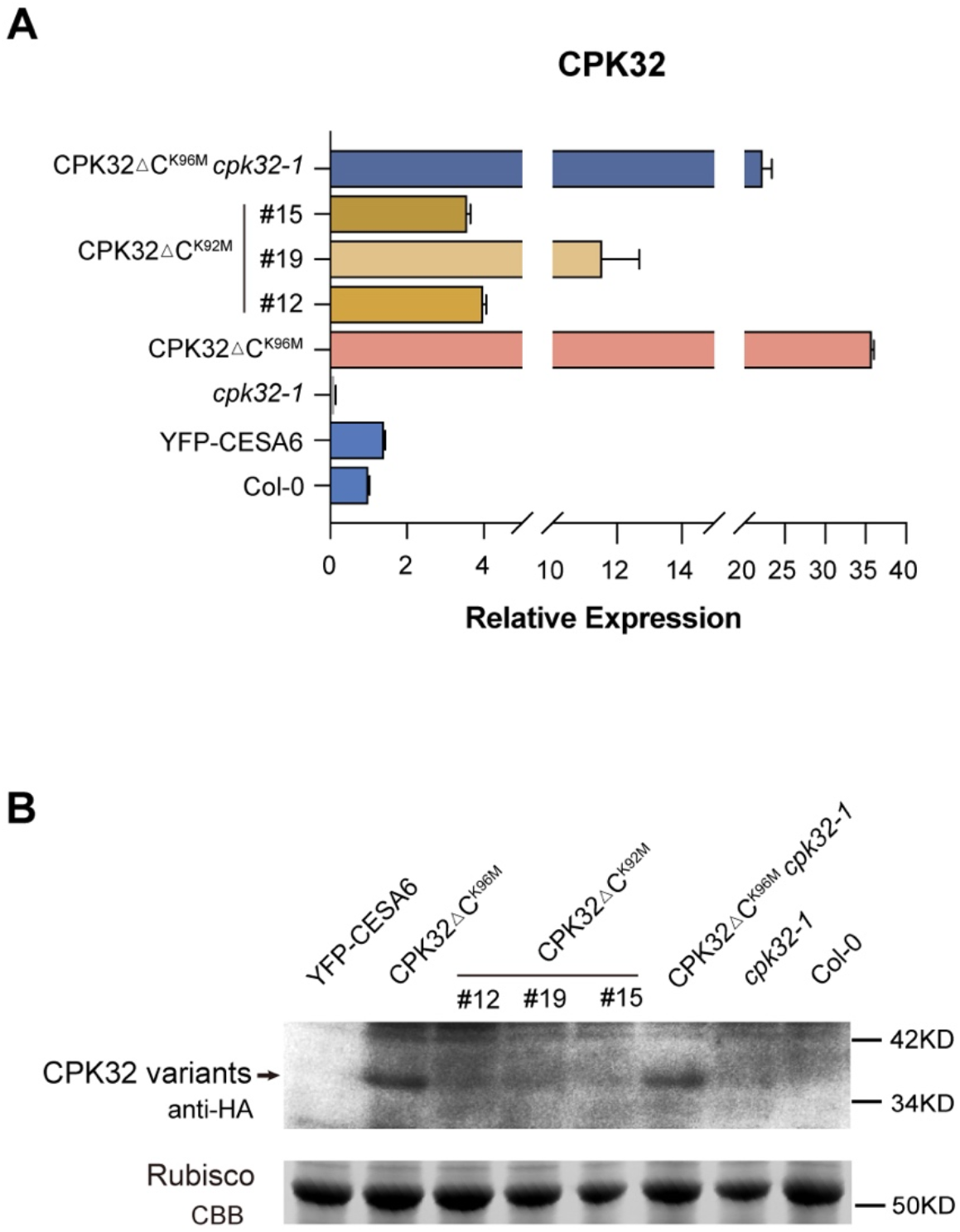
The CPK32ΔC^K92M^ protein was degraded *in vivo*. (A) Histograms indicate the relative expression of *CPK32* in Col-0, *CPK32ΔC*, *CPK32ΔC^K96M^*, *CPK32ΔC^K96M^ cpk32-1*, *YFP-CESA6 prc1-1* and *CPK32ΔC^K92M^*,adjusted by the expression of *ACT2*, normalized to the expression of *CPK32* in Col-0, assessed by RT-qPCR. (B) Western blot analysis of HA-CPK32 protein in Col-0, *cpk32-1*, *CPK32ΔC^K96M^*, *CPK32ΔC^K96M^ cpk32-1*, *YFP-CESA6 prc1-1* and *CPK32ΔC^K92M^*. Upper panel show the immunoblot and lower panel is the loading control.

**Supplementary Fig. S5.**
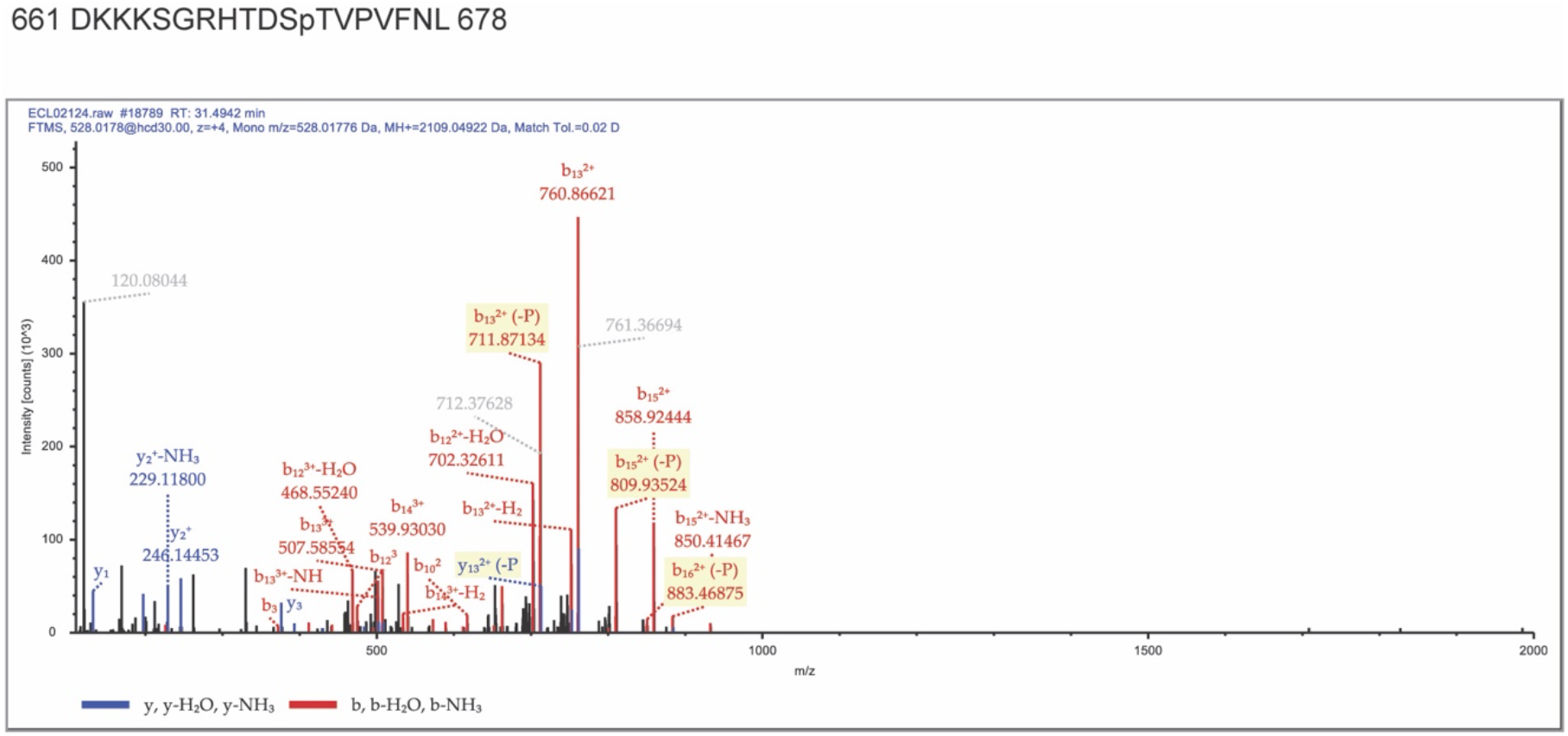
Identification of phosphorylation site of CESA3CD via tandem mass spectrometry. The phosphorylated proteins were separated by SDS-PAGE. The protein bands between 72 KD and 52 KD were cut for in-gel digestion with trypsin. The resulting peptides were analyzed by LC-MS/MS analyses. The tandem mass spectrometry spectra show single phosphorylation site of peptide D^661^-L^678^. The experimental setting is identical to what was shown in Fig. 2A, except that regular ATP was used in this assay.

**Supplementary Fig. S6.**
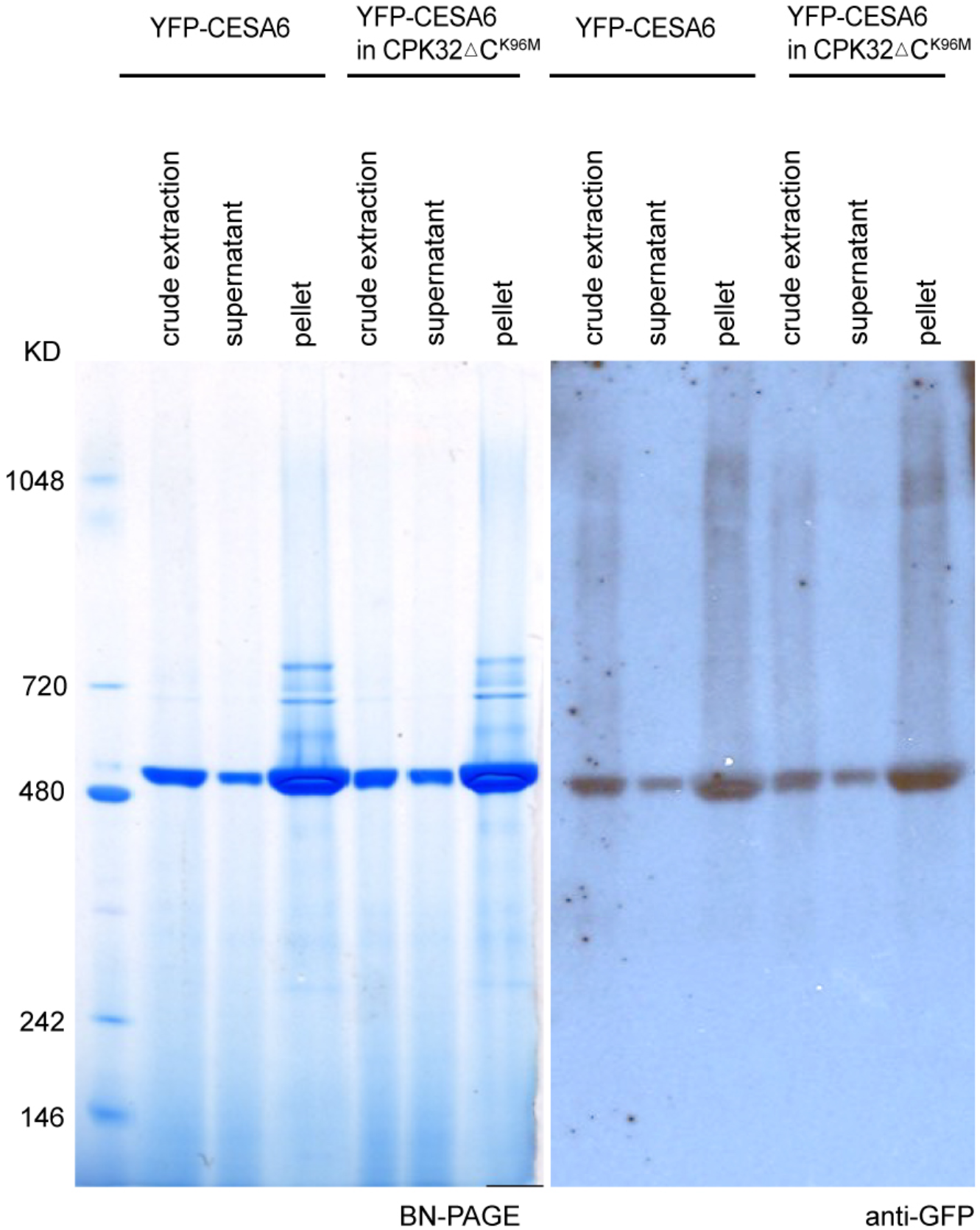
The assembly of CSCs is not affected in *CPK32ΔC^K96M^* seedlings. Western blot analysis of native YFP-CESA6 proteins in crude extraction, supernatant after ultracentrifuge, and pellet containing the microsomal fraction of seedlings in control (YFP-CESA6 mCherry-TUA5) and CPK32ΔC^K96M^ in the same background (YFP-CESA6 mCherry-TUA5). Left panel shows blue-native polyacrylamide gel electrophoresis (BN-PAGE). Right panel shows immunoblot using GFP antibody.

**Supplementary Table 1.**
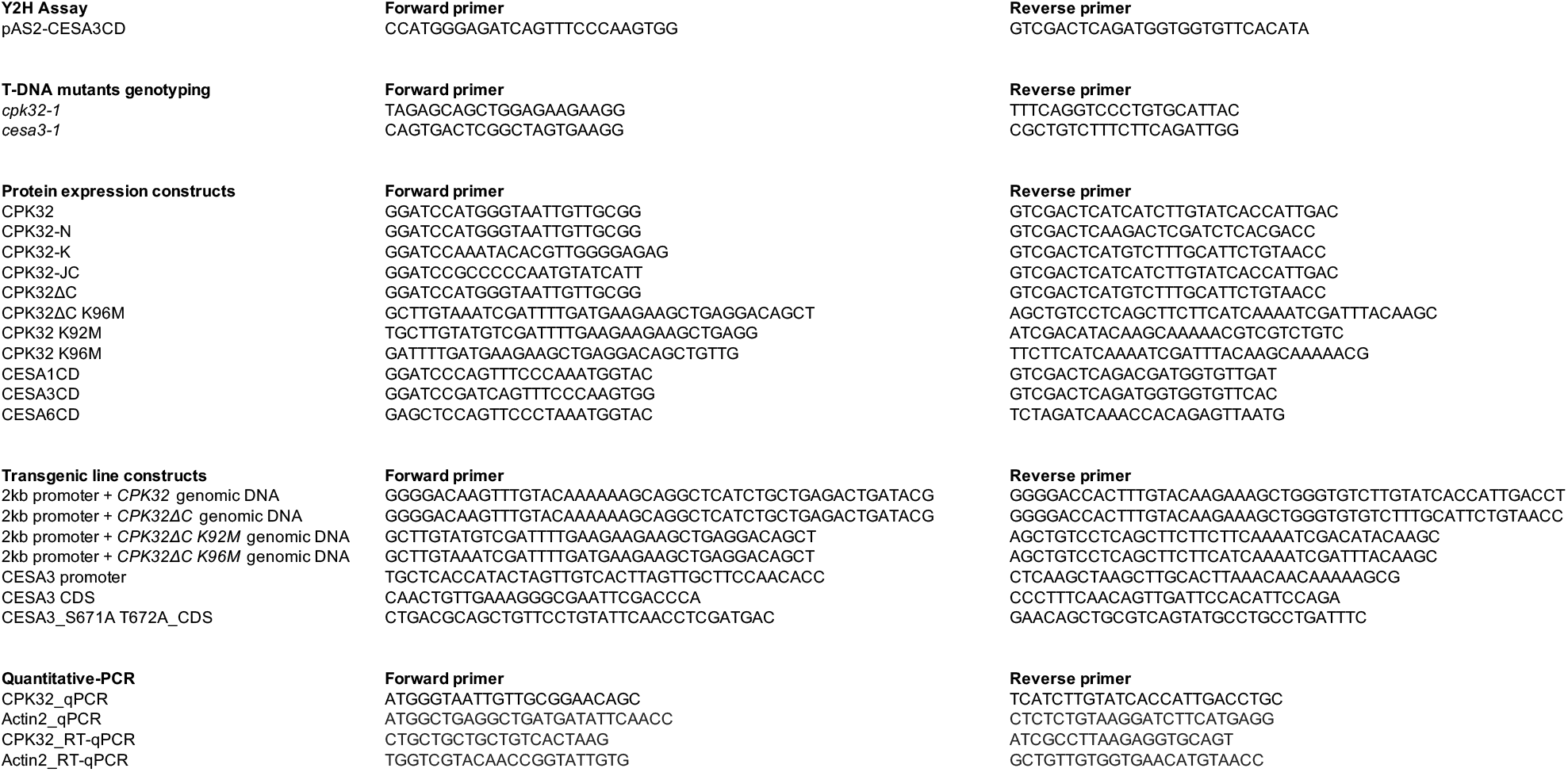
DNA primers used in this study.

