## Supplemental Table 1 for "CALCIUM-DEPENDENT PROTEIN KINASE32 regulates cellulose biosynthesis through post-translational modification of cellulose synthase"

| **Y2H Assay** | **Forward primer** | **Reverse primer** |
|---|---|---|
| pAS2-CESA3CD | CCATGGGAGATCAGTTTCCCAAGTGG | GTCGACTCAGATGGTGGTGTTCACATA |

| **T-DNA mutants genotyping** | **Forward primer** | **Reverse primer** |
|---|---|---|
| *cpk32-1* | TAGAGCAGCTGGAGAAGAAGG | TTTCAGGTCCCTGTGCATTAC |
| *cesa3-1* | CAGTGACTCGGCTAGTGAAGG | CGCTGTCTTTCTTCAGATTGG |

| **Protein expression constructs** | **Forward primer** | **Reverse primer** |
|---|---|---|
| CPK32 | GGATCCATGGGTAATTGTTGCGG | GTCGACTCATCATCTTGTATCACCATTGAC |
| CPK32-N | GGATCCATGGGTAATTGTTGCGG | GTCGACTCAAGACTCGATCTCACGACC |
| CPK32-K | GGATCCAAATACACGTTGGGGAGAG | GTCGACTCATGTCTTTGCATTCTGTAACC |
| CPK32-JC | GGATCCGCCCCCAATGTATCATT | GTCGACTCATCATCTTGTATCACCATTGAC |
| CPK32ΔC | GGATCCATGGGTAATTGTTGCGG | GTCGACTCATGTCTTTGCATTCTGTAACC |
| CPK32ΔC K96M | GCTTGTAAATCGATTTTGATGAAGAAGCTGAGGACAGCT | AGCTGTCCTCAGCTTCTTCATCAAAATCGATTTACAAGC |
| CPK32 K92M | TGCTTGTATGTCGATTTTGAAGAAGAAGCTGAGG | ATCGACATACAAGCAAAAACGTCGTCTGTC |
| CPK32 K96M | GATTTTGATGAAGAAGCTGAGGACAGCTGTTG | TTCTTCATCAAAATCGATTTACAAGCAAAAACG |
| CESA1CD | GGATCCCAGTTTCCCAAATGGTAC | GTCGACTCAGACGATGGTGTTGAT |
| CESA3CD | GGATCCGATCAGTTTCCCAAGTGG | GTCGACTCAGATGGTGGTGTTCAC |
| CESA6CD | GAGCTCCAGTTCCCTAAATGGTAC | TCTAGATCAAACCACAGAGTTAATG |

| **Transgenic line constructs** | **Forward primer** | **Reverse primer** |
|---|---|---|
| 2kb promoter + *CPK32* genomic DNA | GGGGACAAGTTTGTACAAAAAAGCAGGCTCATCTGCTGAGACTGATACG | GGGGACCACTTTGTACAAGAAAGCTGGGTGTCTTGTATCACCATTGACCT |
| 2kb promoter + *CPK32ΔC* genomic DNA | GGGGACAAGTTTGTACAAAAAAGCAGGCTCATCTGCTGAGACTGATACG | GGGGACCACTTTGTACAAGAAAGCTGGGTGTGTCTTTGCATTCTGTAACC |
| 2kb promoter + *CPK32ΔC K92M* genomic DNA | GCTTGTATGTCGATTTTGAAGAAGAAGCTGAGGACAGCT | AGCTGTCCTCAGCTTCTTCTTCAAAATCGACATACAAGC |
| 2kb promoter + *CPK32ΔC K96M* genomic DNA | GCTTGTAAATCGATTTTGATGAAGAAGCTGAGGACAGCT | AGCTGTCCTCAGCTTCTTCATCAAAATCGATTTACAAGC |
| CESA3 promoter | TGCTCACCATACTAGTTGTCACTTAGTTGCTTCCAACACC | CTCAAGCTAAGCTTGCACTTAAACAACAAAAAGCG |
| CESA3 CDS | CAACTGTTGAAAGGGCGAATTCGACCCA | CCCTTTCAACAGTTGATTCCACATTCCAGA |
| CESA3_S671A T672A_CDS | CTGACGCAGCTGTTCCTGTATTCAACCTCGATGAC | GAACAGCTGCGTCAGTATGCCTGCCTGATTTC |

| **Quantitative-PCR** | **Forward primer** | **Reverse primer** |
|---|---|---|
| CPK32_qPCR | ATGGGTAATTGTTGCGGAACAGC | TCATCTTGTATCACCATTGACCTGC |
| Actin2_qPCR | ATGGCTGAGGCTGATGATATTCAACC | CTCTCTGTAAGGATCTTCATGAGG |
| CPK32_RT-qPCR | CTGCTGCTGCTGTCACTAAG | ATCGCCTTAAGAGGTGCAGT |
| Actin2_RT-qPCR | TGGTCGTACAACCGGTATTGTG | GCTGTTGTGGTGAACATGTAACC |
